# Aβ/APP-induced hyperexcitability and dysregulation of homeostatic synaptic plasticity in models of Alzheimer’s disease

**DOI:** 10.1101/2022.01.25.477711

**Authors:** I Martinsson, L Quintino, MG Garcia, SC Konings, L Torres-Garcia, A Svanbergson, O Stange, R England, T Deierborg, JY Li, C Lundberg, GK Gouras

## Abstract

The proper function of the nervous system is dependent on the appropriate timing of neuronal firing. Synapses continually undergo rapid activity-dependent modifications that require feedback mechanisms to maintain network activity within a window in which communication is energy efficient and meaningful. Homeostatic synaptic plasticity (HSP) and homeostatic intrinsic plasticity (HIP) are such negative feedback mechanisms. Accumulating evidence implicates that Alzheimer’s disease (AD)-related amyloid precursor protein (APP) and its cleavage product amyloid-beta (Aβ) play a role in the regulation of neuronal network activity, and in particular HSP. AD features impaired neuronal activity with regional early hyper-activity and Aβ-dependent hyperexcitability has also been demonstrated in AD transgenic mice. We demonstrate similar hyper-activity in AD transgenic neurons in culture that have elevated levels of both human APP and Aβ. To examine the individual roles of APP and Aβ in promoting hyperexcitability we used an APP construct that does not generate Aβ, or elevated Aβ levels independently of APP. Increasing either APP or Aβ in wild type (WT) neurons leads to increased frequency and amplitude of calcium transients. Since HSP/HIP mechanisms normally maintain a setpoint of activity, we examined whether homeostatic synaptic/intrinsic plasticity was altered in AD transgenic neurons. Using methods known to induce HSP/HIP, we demonstrate that APP protein levels are regulated by chronic modulation of activity and show that AD transgenic neurons have an impaired response to global changes in activity. Further, AD transgenic compared to WT neurons failed to adjust the length of their axon initial segments (AIS), an adaptation known to alter excitability. Thus, we present evidence that both APP and Aβ influence neuronal activity and that mechanisms of HSP/HIP are disrupted in neuronal models of AD.

## Background

Alzheimer’s disease (AD) is the leading cause of dementia and the most common neurodegenerative disease. It is characterized by the progressive, age-related accumulation and aggregation of disease-associated proteins, including amyloid-beta (Aβ) that is cleaved from the amyloid precursor protein (APP). This process is thought to drive the loss of synapses and neurons. However, preceding the massive neurodegeneration, AD features aberrant regional neuronal activity in the form of both hyper- and hypo-excitability, and evidence supports that the occurrence of network hyper-excitability early in the disease process is tied to elevated Aβ levels (Vossel *et al*., 2013; Zott *et al*., 2019). However, the precise mechanisms behind this early Aβ-induced hyper-excitability remain unclear. In addition, the normal roles of Aβ/APP in brain physiology and their roles in pathophysiology during AD remain incompletely understood.

The coordinated firing of neurons across networks is considered to be crucial for cognitive function. To maintain their proper function and levels of activity, neurons employ homeostatic synaptic plasticity (HSP) and homeostatic intrinsic plasticity (HIP), which are means by which neurons can tune their activity to the global tonus of activity. Homeostatic scaling is an example of one such tuning mechanism (Turrigiano *et al*., 1998), and other mechanisms of regulation are being investigated. These modulatory processes enable neuronal communication to be maintained within an appropriate window, allowing meaningful information transfer (Turrigiano, 2012). Recently, both APP and Aβ were implicated in the regulation of HSP (Gilbert *et al*., 2016; Galanis *et al*., 2021), demonstrating that these proteins play important roles beyond AD pathophysiology. Further, prior in vivo work in AD transgenic mice suggests that HSP mechanisms might be impaired, since chronic hypo-activity or hyper-activity via either long term sleep deprivation or induction, or unilateral whisker removal, conditions in which HSP should be engaged, negatively impacted AD transgenic compared to wild-type (WT) mice (Kang *et al*., 2009; Tampellini *et al*., 2010).

In this study, we set out to further elucidate the effects of APP and Aβ on neuronal activity. To that end, we utilized primary neuronal cultures from APP/PS1 AD transgenic mice and their WT counterparts and live-cell calcium imaging to perform activity analyses of neuronal networks. We demonstrate that a general increase in transient calcium frequencies occurs in the context of elevated APP and Aβ. Furthermore, we show that specifically CaMKII-positive excitatory neurons from AD transgenic mice exhibit higher amplitude calcium transients. Finally, we demonstrate the impaired ability of AD transgenic compared to WT neurons to properly initiate HSP and HIP mechanisms to adapt to global activity changes.

## Materials and Methods

### Antibodies

The antibodies employed in this study were the following: mouse anti-beta-actin (Sigma-Aldrich, Sweden), rabbit anti-OC against high molecular weight Aβ (Merck Millipore, Sweden), mouse anti-6E10 for human Aβ/APP (BioLegend, Sweden), APPY188 rabbit anti C-terminal APP (Abcam, Sweden), rabbit anti-somatostatin (Abbexa, UK), mouse anti-GAD67 (Merck Millipore, Sweden), mouse anti-CaMKII (Merck Millipore, Sweden), mouseanti-ankyrin-G (Thermo Scientific, Sweden), guinea pig anti-Vglut1 (Synaptic Systems, Germany), rabbit anti-VGAT (Synaptic systems, Germany), rabbit anti-Gephyrin (Synaptic Systems, Germany), and chicken anti-MAP2 (Abcam, UK).

### Neuronal cell culture

Primary neurons were cultured from the cortices and hippocampi of APP/PS1 AD transgenic mouse (APPswe, PSEN1dE9)85Dbo/Mmjax; Jackson Labs, Maine, USA) and APP KO (Jackson labs, Maine, USA, JAX 004133) mouse embryos at embryonic day 15-17 (E15-17). Neurons were cultured as previously described (Willén et al. 2017). Briefly, pregnant mice were anesthetized using isoflurane (MSD Animal Health, Sweden) and sacrificed. Embryos were quickly removed, and biopsies were taken for genotyping. Brains were dissected under constant cooling with chilled (~4 °C) Hanks balanced salt solution (HBSS; Thermo Scientific, Sweden) supplemented with 0.45% glucose (Thermo Scientific, Sweden). Cortices and hippocampi were retrieved and incubated in 0.25% trypsin (Thermo Scientific, Sweden), followed by 2 washes with HBSS. Brain tissue was then triturated in 10% fetal bovine serum (FBS) supplemented Dulbecco’s modified Eagle medium (DMEM; Thermo Scientific, Sweden) with 1% penicillin-streptomycin (Thermo Scientific, Sweden) using glass pipettes until neurons were dissociated. Neurons were plated onto 8 well-plates (for calcium imaging; Ibidi), 6 well plates (for Western blot; Sarstedt, Germany) or glass coverslips in 24 well plates (for immunolabeling; Sarstedt, Germany) coated with Poly-D-lysine (Sigma-Aldrich, Sweden). Neurons were plated with 10% FBS and 1% penicillin-streptomycin in DMEM; following 3-5 h incubation, media was exchanged for complete Neurobasal solution, consisting of Neurobasal medium, B27 supplement, penicillin-streptomycin, and L-glutamine (Thermo Scientific, Sweden). One embryo corresponds to one set of cultures. All animal experiments were performed in accordance with the ethical guidelines and were approved by the Animal Ethical Committee at Lund University ethical permit number 5.8.18-05983/2019.

### Genotyping

Genotyping was carried out using the PCRbio Rapid Extract PCR kit (Techtum, Sweden). In brief, biopsies were incubated with 70 μl distilled H2O, 20 μl 5x PCRbio buffer A (lysis buffer) and 10 μl 10x PCRbio buffer B (protease containing buffer) per vial at 75 °C for 5 min, followed by heating to 95 °C for 10 min. The vials were placed on ice and allowed to cool before vortexing for 3-4 s and centrifuged at 10,000 rpm for 1 min to pellet the debris. The DNA supernatant was then transferred to a new vial. The DNA supernatant was either used directly or stored at −20 °C. For PCR, 1 μl of DNA was incubated with 9.5 μl distilled H_2_O, 12.5 μl 2x PCRbio rapid PCR mix (containing Taq polymerase for DNA amplification), 1 μl primer-set F (APP knockout) and 1 μl primer-set G (APP WT; both 10 μM) for 3 min at 95 °C. The temperature was decreased to 55 °C for 15 seconds to allow for the annealing of primers. The temperature was then increased to 72 °C for 5 min to allow for the extension of DNA. DNA bands were detected using agarose gel electrophoresis.

### Viral vectors

We used lentiviral vectors carrying TdTomato, hAPPwt or hAPPmv (mutant APP resistant to BACE cleavage) under a CaMKII promoter; the genes were inserted via Gene synthesis (Thermo Fischer Scientific) into a plasmid compatible with Gateway technology to serve as an entry clone. Production and titration were performed as previously described (Quintino *et al*., 2013). Primary neurons were transduced at 12-13 days in vitro (DIV) at a multiplicity of infection (MOI) of 5 and analyzed at 19-21 DIV.

### Treatments

Cultured neurons at 19-21 DIV were treated with different compounds before live cell imaging, immunofluorescence or western blot experiments: thiorphan (500 nM, 1 h; Sigma-Aldrich, Sweden), TTX (1 μM, acute (0-1h) or 48 h), bicuculline (20 μM, acute(0-1h) or 48 h), picrotoxin (10 μM, 1 h), CNQX (10 μM, 1 h) (Sigma), and synthetic Aβ1-42 (Tocris, UK) and synthetic reverse Aβ42-1 (Tocris, UK) reconstituted in dimethyl sulfoxide (DMSO) to 250 mM, sonicated 10 min and then centrifuged at 10 000g for 15 min before adding the supernatant to cell culture media.

### Western blot

Cell lysates were prepared using modified RIPA buffer containing 50 mM Tris-HCl (pH 7.4), 150 mM NaCl, 1 mM EGTA, 1% NP-40, 0.25% sodium deoxycholate with added protease and phosphatase inhibitor cocktail II (Sigma-Aldrich, Sweden). BCA protein assay kit (Thermo Scientific, Sweden) was used to determine protein concentrations. Equal amounts of protein from each sample were loaded into 10-20% Tricine sodium dodecyl sulphate– polyacrylamide gel electrophoresis (SDS-PAGE; Sigma-Aldrich, Sweden), followed by immunoblotting on polyvinylidene difluoride (PVDF) membranes (Sigma-Aldrich, Sweden) and intensity quantification was carried out using Image Lab 5.2.1.

### Live-cell imaging

Cultured neurons at 19-21 DIV were incubated with 3 μM of the calcium dye Fluo-4 AM (Thermo Scientific, Sweden) in DMSO(Sigma-Aldrich, Sweden) for 30 min before imaging. Cells were imaged under a Nikon Eclipse Ti microscope at 10 x with 1.4 NA. Live cell imaging chamber (Okolab) was kept at 5% CO2 and 37 °C. Cells were imaged every 100 ms for a duration of 2 min with an iXon Ultra CCD camera (ANDOR Technology).

### Calcium imaging analysis

Time-stacks of calcium imaging files were opened in FiJi; individual Regions of interest (ROIs) were drawn around cell bodies and ROIs were determined to be CaMKII+ or CaMKII-based on TdTomato labeling. Fluorescence intensity over time was extracted, processed and normalized in the MatLab script PeakCaller(Artimovich *et al*., 2017a; b). Spike detection threshold was set to 10% above baseline; for calculation of amplitude heights and interspike intervals; silent neurons were omitted as these would bias the measurement and underestimate the amplitude heights. Spike frequencies and amplitudes were extracted, and raster plots were generated in MatLab.

### Immunofluorescence

Cultured neurons at 19-21 DIV were fixed in 4% paraformaldehyde (PFA) in PBS with 0.12 M sucrose for 20 min, at room temperature (RT). Cells were then blocked in 0.1% saponin (Sigma-Aldrich, Sweden), 1% bovine serum albumin (BSA; Sigma-Aldrich, Sweden) and 2% normal goat serum (NGS; Thermo Scientific, Sweden) in PBS for 1 h at RT. Cells were incubated in primary antibody (diluted in 2% NGS in PBS) overnight at 4 °C. Cells were rinsed in PBS and incubated with secondary antibodies diluted in 2% NGS in PBS. Cells were rinsed in PBS and counterstained with DAPI diluted at 1:2000 (Sigma-Aldrich, Sweden). Imaging was performed with an inverted Olympus IX70 epifluorescence or an inverted Leica SP8 confocal microscope.

### Image analysis

Neurons were labeled for inhibitory or excitatory pre- and post-synaptic markers; CaMKII and VGLUT1 for excitatory synapses and Gephyrin and VGAT for inhibitory synapses. Images were then processed in the ImageJ plugin: SynaptcountJ. SynaptCountJ is a semi-automated plugin for measuring synapse density (Mata *et al*., 2017). By colocalization of two different excitatory or inhibitory synaptic markers one can count the number of excitatory and inhibitory synapses per neuron along with other morphological parameters such as dendritic length. For CaMKII cell quantification 19-21 DIV APP/PS1 and WT neuronal cultures were labeled with DAPI, CaMKII and MAP2. Images were sampled at 20x in an inverted Olympus IX70 epifluorescence microscope and analyzed with the “cell-counter” plugin in ImageJ as percent CaMKII positive out of all MAP2 positive neurons. For analysis of axon initial segment length, ankyrin-G positive axon initial segments were traced and measured in ImageJ by a blinded experimenter.

### Experimental design and statistical analysis

All statistical analyses were performed with GraphPad Prism 8.3. Sample size was denoted as n = number of cells analyzed and N = sets of cultures. Data was first tested for normality using D’Agostino-Pearson omnibus K2 normality test to determine the appropriate statistical test. Mann-Whitney or Kruskal Wallis tests were used to compare distribution of data between groups unless otherwise stated. Correction for multiple testing was performed with Dunn’s correction unless otherwise stated. Graphs are expressed as mean **±** 95% confidence interval with individual values plotted as dots unless otherwise stated in figure legend. Differences were considered significant at *p < 0.05, **p < 0.01, ***p <0.001; ****p<0.0001, n.s., not significant.

## Results

### Increased calcium oscillations in APP/PS1 AD transgenic mouse neurons

To study AD-related neuronal activity alterations *in vitro*, primary cortico-hippocampal neurons from APP/PS1 AD transgenic mice and their WT littermates were loaded with the calcium indicator Fluo-4 AM and imaged for calcium transients using a live cell imaging microscope. Representative raster plots and calcium traces showed that neurons from both APP/PS1 transgenic and WT mice were spontaneously active (Supplemental figure 1). APP/PS1 neurons overall had an increased frequency of calcium transients and higher amplitude of calcium spikes compared to WT neurons (Fig. 1a,b). Inter-spike intervals were also altered in APP/PS1 neurons, which had shorter inter-spike intervals than WT neurons (Fig. 1c). Moreover, APP/PS1 cultures had fewer inactive neurons compared to WT cultures (Fig. 1d), consistent with the other signs of increased excitability.

**Figure 1.**
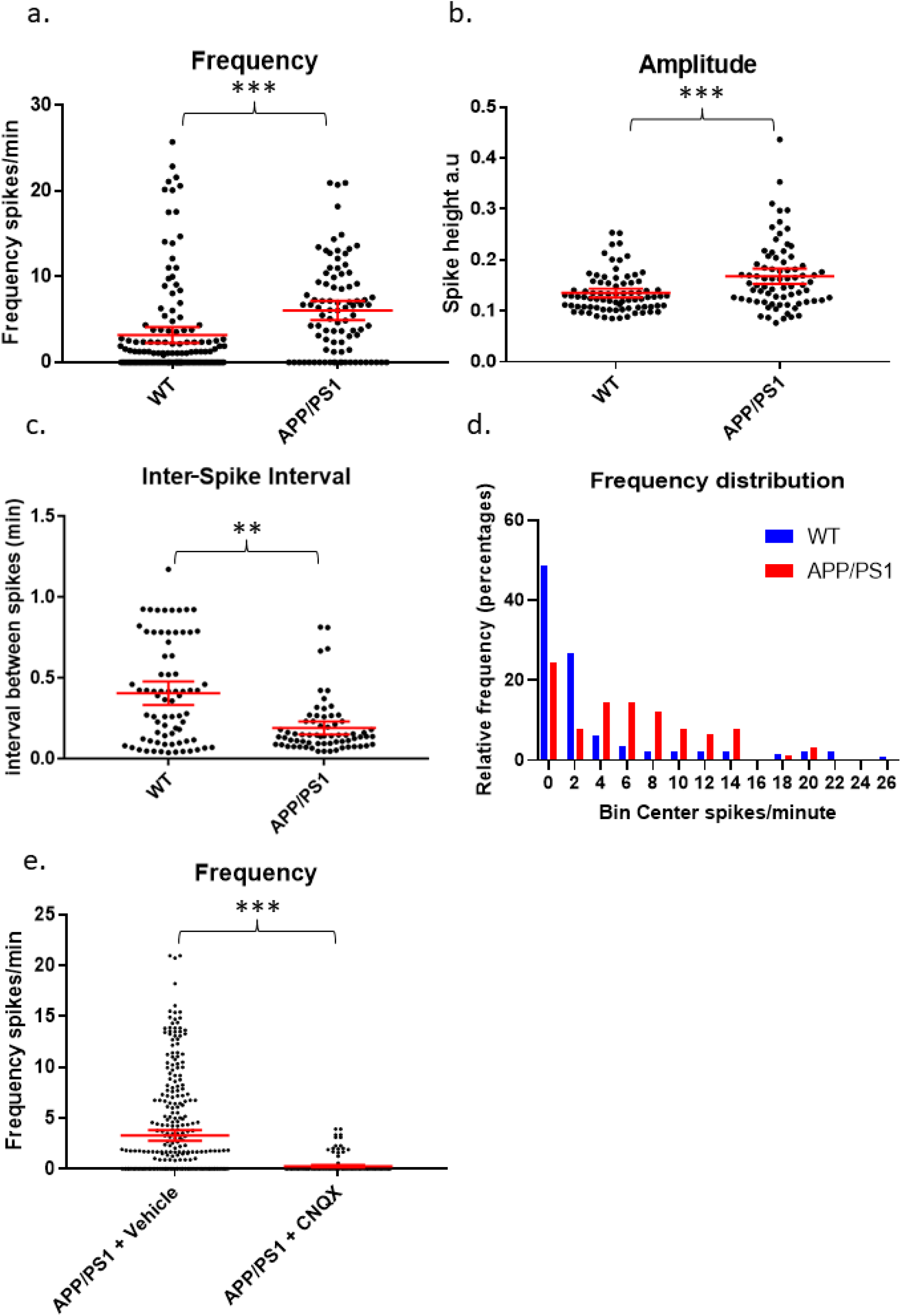
Increased spontaneous activity in APP/PS1 primary neurons. **a)** Frequency of firing (spikes per minute) is increased in APP/PS1 compared to WT neurons (APP/PS1 mean = 6.072, CI = 4.948-7.197, n = 90 compared to WT mean = 3.184, CI = 2.25-4.118, n = 142, p < 0.0001). **b)** Amplitudes of spikes are increased in APP/PS1 compared to WT neurons (APP/PS1 mean = 0.1682, CI = 0.1532-0.1833, n = 76, compared to WT mean = 0.1352, CI = 0.1266-0.1437, n = 80, p = 0.0003). **c)** Inter-spike interval distributions differ between APP/PS1 compared to WT neurons with more of the APP/PS1 neurons having low inter-spike intervals (p = 0.0011 using Kolmogorov-Smirnoff test). **d)** Frequency distribution of firing frequencies of APP/PS1 compared to WT neurons shown in a graph; note higher percentage of WT neurons in the inactive bins; N = 4 cultures. **e)** Graph depicting decrease of firing frequency in APP/PS1 neurons treated with AMPA receptor antagonist CNQX (10 μM; vehicle mean = 3.276, CI = 2.758-3.794, n = 329 compared to CNQX mean = 0.2423, CI = 0.1265-0.3581, n= 172, p = 0.0001); N = 3. Kruskal-Wallis test with Dunn’s correction for multiple comparisons; * p < 0.05, **, p < 0.01 ***, p < 0.001.

As glutamate is the main excitatory neurotransmitter, we next assessed whether glutamate signaling was involved in APP/PS1 neuronal hyperactivity. To do this, APP/PS1 cultures were treated with CNQX to block AMPA receptor-mediated signaling. Following treatment with CNQX, APP/PS1 neurons had a sharp decrease in calcium oscillation frequency and amplitude compared to neurons treated with vehicle (Fig. 1e), supporting the conclusion that glutamate signaling contributes to the hyper-activity of the AD transgenic neuronal cultures.

### Maintained inhibitory GABA signaling and network-level hyperactivity in APP/PS1 neuronal cultures

During early development, GABA can have excitatory effects in culture (Doshina *et al*., 2017). Normally, GABA signaling shifts to being primarily inhibitory at around 14 days *in vitro* (DIV). However, because APP/PS1 neurons may have altered neuronal development due to constitutively over-expressing mutant APP and presenilin (Handler *et al*., 2000; Rama *et al*., 2012), the increased activity seen in 19-21 DIV cultures could, in part, have been due to excitatory GABA signaling. Thus, we tested whether GABA signaling had inhibitory or excitatory effects in our presumably mature 19-21 DIV APP/PS1 neurons by treating them with GABAA blockers bicuculline and picrotoxin. Blocking GABA signaling in the APP/PS1 cultures led to an increased frequency and synchronicity of firing, indicating that GABA had inhibitory effects in our cultures similar to that of WT cultures (Supplemental figure 2a-c).

To further understand what was driving the hyperactivity in APP/PS1 neurons, we examined whether the hyperactivity was being driven at a network or cellular level. To that end, we treated WT and APP/PS1 cultures with tetrodotoxin (TTX), which reduces global activity by blocking sodium channels. The addition of 1 μm TTX stopped most calcium transients in both WT and APP/PS1 neurons, indicating that a global response was achieved and that the hyperactivity in APP/PS1 neurons was at the network-level rather than a cell-specific effect (Supplemental figure 2d-f).

### Hyperactivity of excitatory neurons in APP/PS1 cultures

To parse out the contributions of excitatory and inhibitory neurons in driving the hyperactivity in APP/PS1 cultures, we transduced neurons with a vector to induce the expression of TdTomato under a CaMKII promoter (Fig. 2a), allowing us to distinguish excitatory neurons from GABAergic interneurons. This allowed for the attribution of Fluo-4 measurements to CaMKII-positive excitatory neurons or to CaMKII-negative neurons, which mostly include inhibitory neurons. Interestingly, we detected increased activity in CaMKII-positive APP/PS1 neurons compared to CaMKII-positive WT neurons. In contrast, activity levels in CaMKII-negative neurons were similar between APP/PS1 and WT cultures (Fig. 2b). Furthermore, CaMKII-positive neurons had higher amplitude calcium transients compared to CaMKII-negative neurons in APP/PS1 but not WT cultures (Fig. 2c).

**Figure 2.**
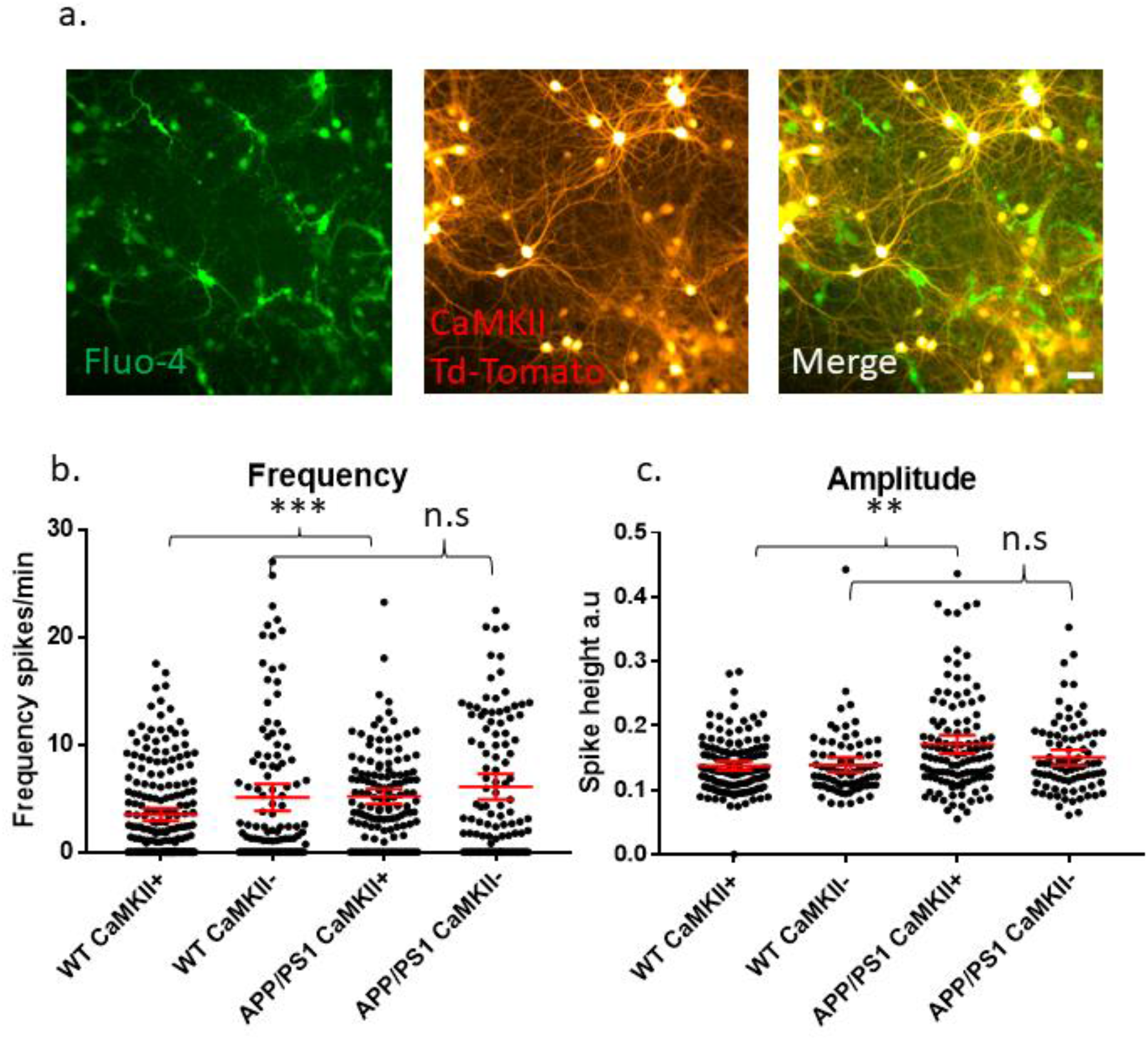
Increased amplitude and frequency of spontaneous calcium transients in CaMKII positive excitatory neurons. **a)** Micrograph provides an example of an image of Fluo-4 (green) and CaMKII TdTomato (red) neurons in culture. **b)** Graph showing increase in spike frequency (spikes per minute) of APP/PS1 CaMKII positive compared to WT CaMKII positive neurons (APP/PS1 mean = 5.176, CI = 4.456-5.896, n = 133 compared to WT mean = 3.479, CI = 2.9-4.057, n = 205, p = 0.0001). In contrast, spike frequency of CaMKII negative APP/PS1 neurons did not significantly differ from WT CaMKII negative neurons (APP/PS1 mean = 6.094, CI = 4.898-7.291, n = 108 compared to WT mean = 5.083, CI = 3.837-6.328, n = 114, p = 0.124). **c)** Amplitude of transients is increased specifically in APP/PS1 compared to WT CaMKII positive excitatory neurons (APP/PS1 mean = 0.171, CI 0.1566-0.1856, n = 115, compared to WT mean = 0.1375, CI = 0.1304-0.1446, n = 138, p = 0.0024). In contrast, the amplitude of transients in CaMKII negative neurons did not differ between APP/PS1 and WT neurons (APP/PS1 mean = 0.1505, CI = 0.1385-0.1625, n = 85, compared to WT mean = 0.1391, CI = 0.1275-0.1506, n = 78, p = 0.302); N = 3. Kruskal-Wallis test with Dunn’s correction for multiple comparisons.; * p < 0.05, ** p < 0.01 *** p < 0.001; scale bar: 50 μm.

We next sought to investigate whether an imbalance in the proportion of excitatory to inhibitory neurons and synapses could underlie the increased levels of activity in APP/PS1 neurons. To do this, we evaluated the relative levels of select proteins known to be localized in either excitatory or inhibitory neurons. Immunoblotting against CaMKII and GAD67, markers expressed by nearly all excitatory and inhibitory neurons, respectively, showed no differences in the levels of CaMKII and GAD67 between APP/PS1 and WT neurons (Fig. 3a-c). Likewise, analyses of neurons immunolabelled for glutamatergic (VGluT and CaMKII) and GABAergic (vGAT and gephyrin) synaptic markers using the image analysis plugins NeuronJ and Synapcount (Mata *et al*., 2017) did not show a significant difference between APP/PS1 and WT cultures (Fig. 3d-f). Similarly, counting CaMKII positive cells per culture did not show a significant difference between the percentage of CaMKII neurons in WT and APP/PS1 neuronal cultures (Fig. 3g). However, consistent with prior work (Siskova *et al*., 2014), our analyses did show decreased dendritic length in APP/PS1 compared to WT neurons (Fig. 3h-i).

**Figure 3.**
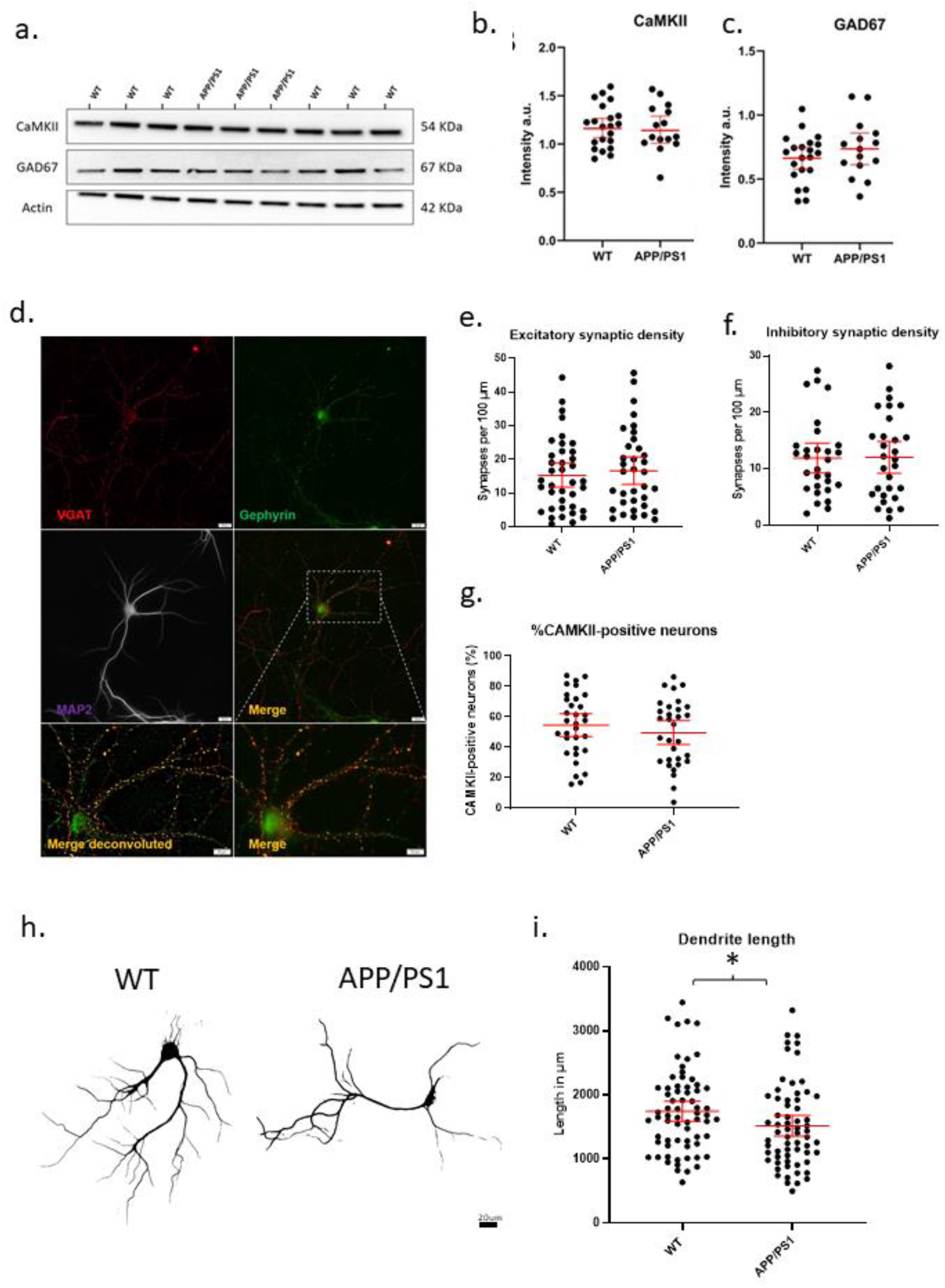
No evidence for gross imbalances in excitatory/inhibitory neurons/synapses in APP/PS1 compared to WT neurons. **a)** Representative western blot of CaMKII and GAD67 protein levels with actin as loading control. **b)** Quantification of western blot of CaMKII in a) (APP/PS1 mean = 1.164, CI = 1.030-1.298, n = 15 and WT mean = 1.181, CI = 1.080-1.282, n = 21, p = 0.8270). **c)** Quantification of western blot of GAD67 in a) (APP/PS1 mean = 0.7359, CI = 0.6129-0.8589, n = 15 and WT mean = 0.6620, CI = 0.5774-0.7466, n = 21, p = 0.2860, unpaired t-test). **d)** Representative micrograph showing WT neuron labeled with VGAT (red), Gephyrin (green) and MAP2 (magenta). Lower panels show an enlarged view of neuron both with (left) and without (right) deconvolution. Scale bars = 20 μm and 10 μm. **e)** Graph depicting excitatory synaptic density from VGLUT/CAMKII overlap divided by neurite length (APP/PS1 mean = 16.64., CI = 12.53-20.74, n = 35 and WT mean = 15.28, CI = 11.74-18.83, n = 38, p = 0.6129, unpaired t-test). **f)** Graph depicting inhibitory synaptic density from VGAT/Gephyrin overlap divided by neurite length (APP/PS1 mean = 12.00, CI = 9.160-14.84, n = 29 and WT mean = 11.87, CI = 9.243-14.50, n = 29 p = 0.9458). **g)** Graph depicting quantification of percentage CaMKII neurons in WT and AD cultures. (APP/PS1 mean = 49.61% CaMKII positive neurons CI = 41.67-57.54% compared to WT mean = 54.64% CI = 47.08-62.14%, p = 0.35, unpaired t-test). **h)** Representative binary images of WT and APP/PS1 neurons labeled for MAP2; scale bar = 20 μm. **i)** Graph showing decreased dendrite length in APP/PS1 compared to WT neurons (APP/PS1 mean = 1518, CI = 1355-1682, n = 64, and WT mean = 1744, CI = 1588-1900. N = 67, p = 0.045, unpaired t-test).

### Individual contributions of APP and Aβ to neuronal hyperactivity

To dissect out the individual role of APP on neuronal activity, we investigated whether APP over-expression alone without a concomitant elevation in Aβ could cause hyper-activity by transducing WT neurons with constructs encoding either mutant human APP resistant to BACE cleavage (hAPPmv)(Kamenetz *et al*., 2003) or WT human APP (hAPPwt), both under a CaMKII promoter. Remarkably, the expression of either APP construct in WT neurons led to a robust increase in the frequency and amplitude of calcium transients (Fig. 4a,b), indicating an Aβ-independent effect of APP on neuronal activity in excitatory neurons.

**Figure 4.**
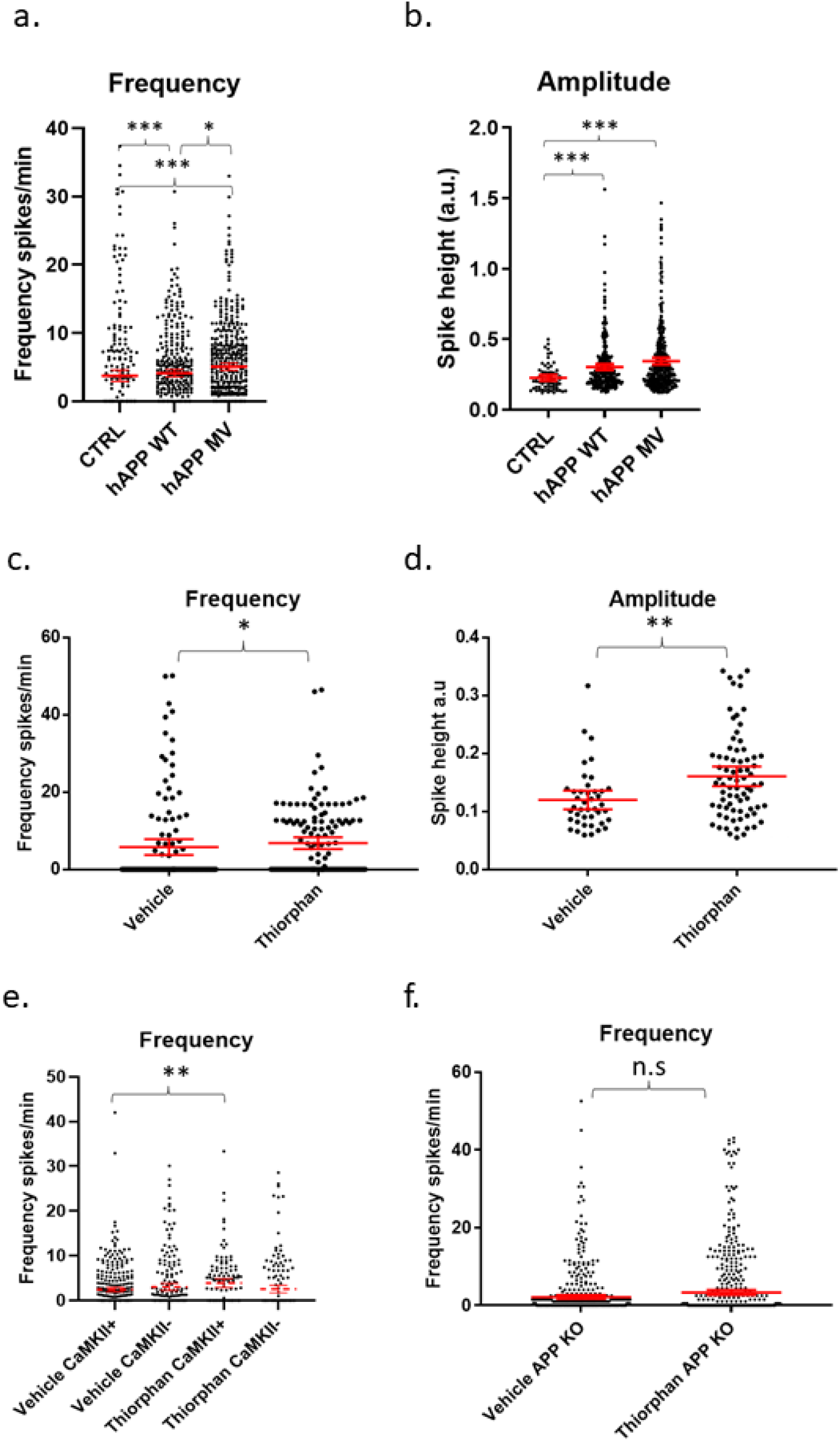
APP and Aβ increase calcium oscillation frequency and amplitude. **a)** Graph depicts firing frequencies in WT neurons transduced with a viral vector harboring hAPPwt or hAPPmv (BACE cleavage resistant) under a CaMKII promoter. Note that both hAPPwt and hAPPmv increase the firing frequency compared to control (CTRL), with hAPPmv having a stronger effect (hAPPmv mean = 5.087, CI = 4.542-5.633, n = 436; hAPPwt mean = 4.112, CI = 3.585-4.640, n = 375 and CTRL mean = 3.757, CI = 2.925-4.588, n = 292, p values respective to CTRL = hAPPmv = 0.0001, hAPPwt = 0.0001, p value hAPPmv compared to hAPPwt (p = 0.0259), Kruskal Wallis Dunn’s correction). **B)** Graph showing increased amplitude in neurons expressing hAPPwt and hAPPmv under CaMKII promoter (hAPPmv mean = 0.3444, CI = 0.3197-0.3691, n = 344 and hAPPwt mean = 0.3021, CI = 0.2808-0.3433, n = 278 and CTRL mean = 0.2256, CI = 0.2034-0.2479, n = 67,, N = 3, p values respective to CTRL; hAPPmv = 0.0001, hAPPwt = 0.0004, p value, Kruskal Wallis Dunn’s correction.). Kruskal-Wallis test with Dunn’s correction for multiple comparisons.; * p < 0.05, ** p < 0.01 *** p < 0.001. **c)** WT neurons treated with 500 nM of the neprilysin inhibitor thiorphan for 1 hour show increased firing frequency (thiorphan mean = 6.814, CI = 5.283-8.344, n = 133, compared to vehicle mean = 5.783, CI = 3.749-7.817, n = 125, N = 3, p = 0.0101). **d)** The graph depicts increased spike amplitudes after 1 hour of treatment of WT neurons with thiorphan (thiorphan mean = 0.1605, CI = 0.1436-0.1774, n = 76, compared to vehicle mean = 0.1198, CI = 0.1036-0.136, n = 42, p = 0.0012). **e)** Graph showing increase in spike frequency of thiorphan treated CaMKII positive compared to vehicle treated CaMKII positive neurons (thiorphan mean = 3.905, CI = 2.942-4.869, n = 128 compared to vehicle mean = 2.432, CI = 1.934-2.931, n = 332, N = 3, p = 0.0089). In contrast, spike frequency of CaMKII negative thiorphan treated neurons did not significantly differ from vehicle treated CaMKII negative neurons (thiorphan mean = 2.566, CI = 1.726-3.406, n = 173 compared to vehicle mean = 3.018, CI = 2.243-3.794, n = 218, p = 0.0894). **f)** Graph showing how thiorphan was ineffective at inducing increased frequency in APP KO neurons compared to vehicle (APP KO thiorphan mean = 3.329, CI = 2.692-3.967, n = 601 compared to APP KO vehicle mean = 2.196, CI = 1.689-2.704, n = 514, N = 3, p = 0.8604, Mann-Whitney U-test).

Likewise, we sought to investigate whether increased Aβ levels alone could induce hyper-activity. We wanted to increase Aβ levels in WT cultures without affecting APP and PS1 as this could confound our results, since we provide evidence that overexpression of APP has Aβ-independent effects on neuronal activity, while PS1 has been shown to alter calcium signaling (Lerdkrai *et al*., 2018). Therefore, we utilized an inhibitor of the Aβ degrading enzyme neprilysin, which increases Aβ levels (Abramov *et al*., 2009), primarily at synapses, as neprilysin is highly expressed pre-synaptically (Iwata *et al*., 2004; Abramov *et al*., 2009). After 1 h of treatment with the neprilysin inhibitor thiorphan (500 nM), calcium transient frequencies (Fig. 4c) and amplitudes (Fig. 4d) were increased in WT primary neurons. Interestingly, thiorphan led to a greater increase in firing frequency in CaMKII-positive compared to CaMKII-negative neurons (Fig. 4e). As neprilysin also degrades other peptides, such as substance P and neurokinin A, as a control, we assessed the effect of thiorphan treatment on APP knockout (KO) neurons, which lack APP and, thus, the capacity to generate Aβ. Indeed, calcium transient frequencies and amplitudes from APP KO neurons treated with thiorphan did not significantly differ from APP KO neurons treated with vehicle alone (Fig. 4f), supporting the conclusion of elevated Aβ levels as driving the hyperactivity in WT neurons treated with thiorphan. However, APP KO neurons have been shown to have altered synaptic composition (Martinsson *et al*., 2019) and calcium transients (Yang *et al*., 2009), which could potentially mask effects by thiorphan. Therefore, as an alternative to investigate elevated Aβ levels and hyperactivity, we added exogenous synthetic Aβ peptide to WT cultures.

Immunolabeling WT neuronal cultures treated with 0.5 μM of synthetic human Aβ1-42 for 2 hours with the human-specific Aβ/APP antibody 6E10 showed that the added exogenous Aβ1-42 localized to the dendritic spines of CaMKII-TdTomato expressing neurons (Fig. 5a, b). This was consistent with prior findings showing that exogenous Aβ1-42 preferentially binds to synaptses of CaMKII-immunoreactive neurons (Willen *et al*., 2017). Interestingly, we detected marked colocalization of the added human Aβ1-42 with the fibril and fibrillar oligomer-specific antibody OC that detects amyloid structures (Hatami *et al*., 2014), consistent with aggregation of the exogenous Aβ1-42 at synapses, which is consistant with prior work (Willén *et al*., 2017). While for these experiments we added supraphysiological levels of Aβ1-42 (0.5 μM), in order to more readily visualize its localization we next assayed what effect more physiological Aβ1-42 increases would have on calcium oscillations. Given that physiological levels of Aβ are in the picomolar range and that picomolar levels of exogenous Aβ were reported to increase LTP (Puzzo *et al*., 2008), we added 200 pM of synthetic human Aβ1-42 acutely to mouse WT neuronal cultures expressing CaMKII-driven TdTomato. Addition of 200 pM synthetic Aβ to WT cultures led to modest increases in the firing frequencies of both CaMKII-positive and CaMKII-negative neurons (Fig. 5c). In addition, these picomolar levels of Aβ1-42 led to increased amplitudes of calcium oscillations in CaMKII-positive neurons but not CaMKII-negative neurons (Fig. 5d). However, higher concentrations of synthetic Aβ1-42 (500 nM) did not lead to significantly increased activity (Fig. 5e), though 500 nM of synthetic Aβ1-40 did lead to a robust increase in firing frequency. In summary, our results support the concept that APP and Aβ can independently induce increases in synaptic activity, which likely plays a role under physiological and pathological conditions.

**Figure 5.**
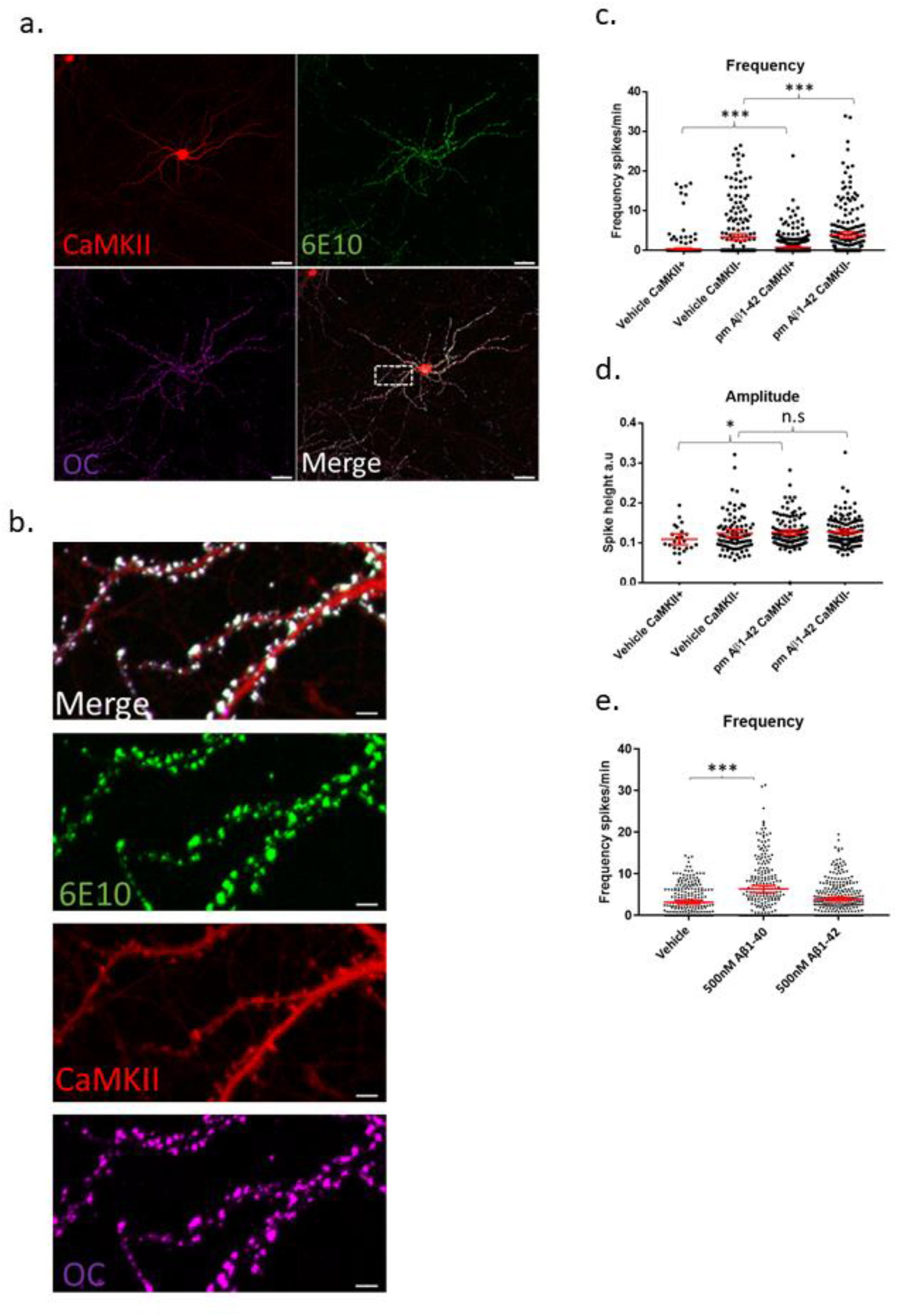
Aβ preferentially binds synaptic compartments on CaMKII positive neurons and appears to have a dose dependent effect on spike frequency. **a)** Micrograph showing that synthetic human Aβ1-42 preferentially binds to dendrites of CaMKII positive murine WT neurons; Td Tomato expressed through CaMKII promoter, human specific anti-Aβ antibody 6E10 (green) and conformation specific anti-amyloid antibody OC (magenta); scale bar = 50 μm. **b)** Insert from a) showing Aβ1-42 targeting synapses; note that antibody 6E10 labels the added synthetic human Aβ and that this is also labeled by the fibrillar oligomer antibody OC; scale bar = 5 μm. **c)** WT neurons treated with 200 pM synthetic Aβ1-42 show increased spike frequency compared to DMSO treated vehicle control neurons both in CaMKII positive neurons (Aβ1-42 mean = 0.7554, CI = 0.5724-0.9383, n = 477, compared to vehicle mean = 0.367, CI = 0.1636-0.5704, n = 401, p < 0.0001) and in CaMKII negative neurons (Aβ1-42 mean = 3.829, CI = 3.033-4.685, n = 214, compared to vehicle mean = 3.346, CI = 2.508-4.183, n = 218). **d)** Graph depicting increased amplitude in CaMKII positive neurons treated with 200 pM Aβ1-42 compared with DMSO vehicle control (Aβ1-42 mean = 0.1256, CI = 0.119-0.1321, n = 128, compared to vehicle mean = 0.1087, CI = 0.09495-0.1225, n = 24, p = 0.045). However, CaMKII negative neurons did not show a significant increase in amplitude with Aβ treatment (Aβ1-42 mean = 0.1272, CI = 0.1209-0.1334, n = 133, compared to vehicle mean = 0.1235, CI = 0.114-0.133, n = 94, p = 0.2103). **e)** Graph shows WT neurons treated with 500 nM Aβ1-42 or 500 nM Aβ1-40. While 500 nM Aβ1-42 leads to a slight but not significantly different distribution of activity, Aβ1-40 strongly increases activity (Aβ1-40 mean = 6.344, CI = 5.503-7.186, n = 235 and Aβ1-42 mean = 3.901, CI = 3.439-4.363, n = 287 compared to vehicle mean = 3.194, CI = 2.752-3.635, n = 255; p = 0.0001 and p = 0.14.); Kruskal Wallis test, N = 3.

### Dysregulated homeostatic plasticity in APP/PS1 neurons

Since we showed that elevating either APP or Aβ in WT neurons can increase neuronal activity, we hypothesized that the continuously high levels of APP and Aβ in APP/PS1 neurons disrupt neuronal network activity and function. We speculated that HSP and HIP mechanisms, which help maintain synaptic firing within the boundaries of meaningful communication (Turrigiano, 2008), may be impaired in AD transgenic neurons and, as a result, may no longer be effective at returning activity levels back to a baseline. Dysfunctional HSP/HIP could thus explain part of the sustained hyperexcitability observed in neurons from AD transgenic mouse models. We therefore hypothesized that long-term high levels of Aβ/APP might impair homeostatic plasticity. This can be tested by manipulating neuronal firing outside of a network’s set point, leading to plasticity changes that maintain the baseline activity level. We therefore initially treated APP/PS1 and WT neurons with TTX and bicuculline, which should immediately decrease network activity and increase network activity, respectively. Indeed, acute treatment of WT and APP/PS1 neurons with TTX and bicuculline led to the expected decreased and increased activity, respectively (Supplementary figure 2e-f). Next we treated WT and APP/PS1 neurons with TTX or bicuculline for a longer period of time (48 hours), which are established methods for inducing HSP/HIP (Turrigiano *et al*., 1998). Interestingly, 48 hours of TTX treatment led to a sharp decrease by western blot in total APP protein levels in both WT and APP/PS1 (Fig. 6a-b), which was consistent with immunolabeling of cultures treated with TTX or bicuculline (Fig. 6c). There was also a trend for a decrease in CaMKIIα, which was previously reported to be downregulated with TTX induced homeostatic scaling (Thiagarajan *et al*., 2002). To determine whether HSP and HIP mechanisms were induced by the long-term treatments, we measured the calcium transients after HSP induction. Treating WT neurons with TTX or bicuculline for 48 hours led to the expected changes in excitability; most TTX treated WT neurons recover their ability to fire and while the distribution of firing rates is altered (Fig. 6d) the mean firing frequency does not differ significantly from WT vehicle treated neurons after 48 hours (Fig. 6f). In response to the 48 hours of treatment with bicuculline, which acutely elevates activity, most WT neurons significantly decreased their firing frequency. Of note, the extended bicuculline treatment in WT neurons appears to lead to two types of firing, with one group of neurons maintaining a high firing frequency and another group that is silent (Fig. 6d,f). In contrast, the firing frequency of APP/PS1 neurons treated for 48 h with bicuculline remained increased, while the firing frequency of APP/PS1 neurons after chronic TTX treatment remained low (Fig. 6e,g). Thus, APP/PS1 neurons did not respond to the prolonged bicuculline or TTX exposure as the WT neurons, indicating that the APP/PS1 neurons were unable to compensate to these pharmacologically-induced changes in neuronal activity.

**Figure 6.**
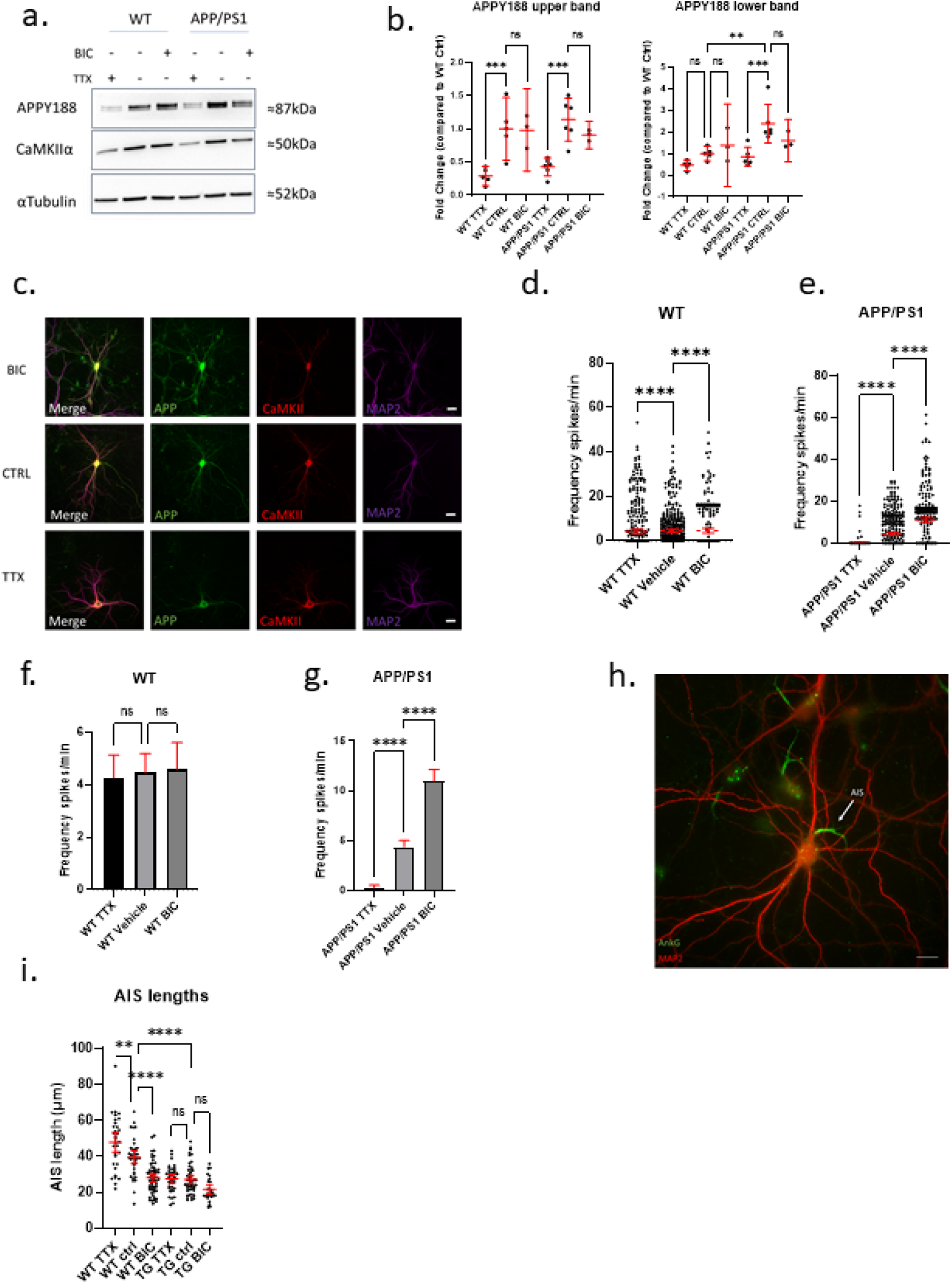
Neurons from AD transgenic mice are unable to initiate homeostatic synaptic/intrinsic plasticity. **a)** Representative western blot of APP and CaMKIIα in WT and APP/PS1 neurons treated with TTX or bicuculline for 48 hours; αTubulin is used as a loading control. **b)** Graph depicts quantification of fold change from western blots in a), APPY188 upper band (WT TTX mean = 0.2865, CI 0.1287-0.4244, n = 5, WT control mean =1.000, CI 0.5298-1.470, n = 5, WT bicuculline mean = 0.9782, CI 0.3574-1.599, n = 3, APP/PS1 TTX mean = 0.4251, CI 0.2893-0.5609, n = 6, APP/PS1 control mean = 1.137, CI 0.8116-1.462, n = 6, APP/PS1 bicuculline mean = 0.9038, CI 0.6975-1.110, n = 3) and APPY188 lower band (WT TTX mean = 0.4605, CI 0.1990-0.7220, n = 5, WT control mean =1.000, CI 0.6601-1.340, n = 5, WT bicuculline mean = 1.386, CI −0.5278-3.299, n = 3, APP/PS1 TTX mean = 0.8590, CI 0.4258-1.292, n = 6, APP/PS1 control mean = 2.239, CI 1.493-3.290, n = 6, APP/PS1 bicuculline mean = 1.602, CI 0.6279-2.576, n = 3). Note that upper APP band is lower in TTX treated WT compared to WT control p = 0.0006 and TTX treated APP/PS1 compared to APP/PS1 control p = 0.0002. For APP lower band APP/PS1 control was significantly higher than WT control p = 0.0017 while APP/PS1 TTX was lower than APP/PS1 control p = 0.0003 (one-way ANOVA, Šidak correction). **c)** Micrograph showing APPY188 and CaMKIIα labelling after 48 hours of TTX or Bicuclline treatment. Note that APPY188 labeling is relatively weaker in TTX treated neuron consistent with the results from western blot. Scale bar = 20 μm. **d)** Graph demonstrating HSP adaptations in WT cultures treated with TTX or bicuculline for 48 hours. Note that the distribution of oscillation frequency in bicuculline treated neurons is differnt compared to vehicle treated (TTX mean = 4.265, CI 3.397-5.134, n = 442; vehicle mean = 4.467, CI = 3.741-5.192, n = 413; bicuculline mean = 4.593, CI = 3.574-5.611, n = 303, p values compared to vehicle: TTX = 0.0001 and bicuculline = 0.0001, Kruskal-Wallis test, N = 4). **e)** Graph showing absence of adaptation in APP/PS1 neurons in response to 48 hours of TTX or bicuculline treatment. Note how both TTX and bicuculline treated APP/PS1 are largely opposite from WT neurons treated with TTX bicuculline after 48 hours. (TTX mean = 0.272, CI 0.0154-0.527, n = 210; vehicle mean = 4.368, CI = 3.729-5.008, n = 464; bicuculline mean = 10.990, CI = 9.835-12.140, n = 368, p values compared to vehicle: TTX = 0.0001 and bicuculline = 0.0001, Kruskal-Wallis test, N = 4). **f)** bar graph showing mean firing frequency and CI from d) note that mean firing frequencies are similar between the groups. (WT TTX vs WT vehicle p = 0.9203, WT bicuculline vs WT vehicle, p = 0.9733, one-way ANOVA, dunnett’s test). **g)** bar graph showing mean firing frequency and CI from e) note that mean firing frequencies are do not recover as in WT. (APP/PS1 TTX vs APP/PS1 vehicle p = 0.0001, APP/PS1 bicuculline vs APP/PS1 vehicle, p = 0.0001, one-way ANOVA, dunnet’s test). **h)** Micrograph depicting WT neuron labeled with MAP2 (red) and ankyrin-G (green), which labels the AIS. White arrow points to AIS; scale bar = 20 μm. **i)** Graph shows quantification of AIS lengths after treatment with TTX or bicuculline for 48 hours (WT TTX mean = 47.76, CI = 42.36-53.17, n = 32; WT vehicle mean = 39.44, CI =35.87-43.02, n = 37; WT bicuculline mean = 28.07, CI = 25.75-30.39, n = 57; APP/PS1 TTX mean = 27.52, CI = 25.01-30.04, n = 35; APP/PS1 vehicle mean = 27.20, CI = 24.85-29.55, n = 49; APP/PS1 bicuculline mean = 21.57, CI = 18.86-24.27, n = 26); ordinary one way ANOVA, Tukey’s multiple comparisons test.

To explore a mechanistic aspect of these HSP/HIP alterations we investigated axon initial segment (AIS) modifications in the chronic TTX and bicuculline treated cultures, as lengthening or shortening of the AIS modifies the excitability of neurons (Hedstrom *et al*. (2008). By immunolabeling for ankyrin-G, a protein involved in linking voltage-gated channels to the AIS, we could identify the AIS and measure its length (Hedstrom *et al*., 2008). In WT neurons, 48 hours of treatment with either bicuculline or TTX led to the expected decreased and increased AIS lengths, respectively (Fig. 6h, i). In contrast, APP/PS1 neurons did not display adjustments of the AIS length upon either of these HSP/HIP-inducing treatments. Interestingly, the AIS lengths of APP/PS1 neurons were already shorter than those of WT neurons at baseline. Together these experiments demonstrate an inability of APP/PS1 neurons to use HSP/HIP mechanisms to adapt to extrinsic changes in activity.

## Discussion

Here we present evidence that homeostatic plasticity mechanisms are disrupted in APP/PS1 AD transgenic neurons. While Aβ and APP have recently been implicated in normal HSP (Gilbert *et al*., 2016; Galanis *et al*., 2021), we present evidence of dysfunctional homeostatic plasticity in AD transgenic neurons. This could help explain why AD transgenic mice are more susceptible to pharmacologically-induced and spontaneous seizures (Minkeviciene *et al*., 2009; Reyes-Marin & Nuñez, 2017) and could provide a framework for explaining the increased sensitivity to seizures in AD patients (Pandis & Scarmeas, 2012). Furthermore, we present data consistent with hyperexcitability in APP/PS1 compared to WT neurons, which we show occurs mainly in excitatory neurons. The calcium oscillation hyper-activity that we observed appeared to be network-driven rather than cell intrinsic. Moreover, part of the hyper-activity in AD transgenic neurons may be caused by the over-expression of APP independent of Aβ. However, elevated levels of Aβ alone also increased activity.

Similar to previous studies, we found that exogenously added Aβ1-42 primarily targeted synapses of CaMKII-positive neurons, specifically in a dendritic rather than axonal pattern (Lacor *et al*., 2004; Willen *et al*., 2017). However, this does not exclude the possibility of Aβ binding to presynaptic terminals proximal to dendrites. Still, the binding of Aβ1-42 to CaMKII-positive synapses might explain why excitatory neurons, in particular, are affected.

The increased frequencies and amplitudes of calcium oscillations that we observed with neprilysin inhibition as a means to elevate endogenous Aβ are consistent with results from Abramov *et al*. (2009), showing increased release probability with thiorphan treatment in cultured hippocampal neurons. However, since neprilysin also degrades various enkephalins and peptide neurotransmitters, such as substance P, which can influence calcium stores (Heath *et al*., 1994), we additionally showed a lack of effect on calcium oscillations with neprilysin inhibition in APP KO neurons.

As the calcium oscillations were increased in APP/PS1 compared to wild type neurons, we also considered whether this might be due to decreased inhibitory interneurons/synapses. However, we did not see significant differences in the protein levels of GAD67 and CaMKII, markers of GABAergic interneurons and excitatory neurons, respectively. Further, we did not detect significant differences in the synaptic density of either excitatory or inhibitory synapses. While neuropathological studies have emphasized the vulnerability of select classes of excitatory projection neurons (Stranahan & Mattson, 2010), the relative involvement of different inhibitory and excitatory neurons in early Aβ-induced hyperactivity remains poorly defined. Increasing evidence suggests that alterations in inhibitory neuron connectivity lead to changes in network functions in AD. For example, increased parvalbumin and gephyrin labeling perisomatically in CA1 neurons of young APP/PS1 transgenic mice was shown (Hollnagel *et al*., 2019), which might represent an adaptation to increased Aβ/APP and, therefore, increased activity in these mice.

The function of neuronal networks is highly dependent on maintaining homeostatic set points and keeping activity within functional windows. Deviations from these set-points lead to network dysfunction. Structural homeostatic synaptic plasticity is known to occur at three main locations: 1. at the post-synapse involving reduced or increased levels of surface receptors (Turrigiano *et al*., 1998); 2. at the axon initial segment (AIS) by either increasing or decreasing its length or by shifting the AIS further out into the axon (Wefelmeyer *et al*., 2016); or 3. at the pre-synapse by modifying how much neurotransmitter is stored in synaptic vesicles or through homeostatic maintenance of presynaptic exocytosis (Delvendahl *et al*., 2019). Of note, Aβ was suggested to overshoot normal homeostatic scaling in response to sensory deprivation *in vivo* or TTX-mediated inhibition *in vitro* (Gilbert *et al*., 2016), and Aβ was recently shown to regulate homeostatic synaptic upscaling after activity blockade in dentate gyrus *in vivo* (Galanis *et al*., 2021). Our findings that APP/PS1 neurons do not increase their activity after 48 hours of TTX treatment nor decrease firing rate after 48 hours of bicuculline treatment suggest that Aβ/APP play an important roles not only in adjusting to activity blockade but also have roles in homeostatic downscaling in response to excessive activity. Interestingly, we found that 48 hours of bicuculline treatment led to a strong reduction in firing in the majority of neurons in WT cultures and another population of neurons that maintained a high firing frequency; a related observation was made in visual cortex homeostatic plasticity following visual deprivation (Barnes *et al*., 2015), where differential adaptations by excitatory and inhibitory neuron populations was described. Interestingly, the mean firing rates are similar between TTX, vehicle and bicuculline treated WT neurons after 48 hours consistent with findings suggesting that while single unit firing is unstable, networks maintain surprisingly stable firing frequencies (Slomowitz *et al*., 2015). We previously showed in primary neurons from Tg2576 AD transgenic mice, which overexpress human APP with the Swedish mutation, reduced AMPA receptor levels in culture (Almeida *et al*., 2005). Whether this is the result of adaptation to higher basal activity levels or a consequence of synaptotoxicity remains to be determined. Moreover, we demonstrated that APP protein levels decrease with chronic TTX treatment. This reduction of APP protein levels could potentially be involved in increasing excitability of excitatory neurons as it has been reported that conditional APP family triple knockout increases excitability of excitatory neurons (Lee *et al*., 2020) and we previously showed that APP knockout increased GluA1 protein levels in cultured neurons (Martinsson *et al*., 2019). While TTX decreased APP, the bicuculline treatment did not significantly alter the APP levels. Neuronal activity can also be modulated by modifying the AIS. We show evidence that the AIS is lengthened with TTX treatment and shortened after treatment with bicuculline in WT neurons, which, however, did not occur in APP/PS1 neurons. Shortening the AIS is a way to decrease intrinsic excitability and previous work in slice cultures has shown that 1 hour of bicuculline treatment was sufficient to decrease the length of the AIS (Jamann *et al*., 2021). Since the average lengths of the AIS in APP/PS1 neurons are shorter at baseline than in WT neurons, and are not altered by chronic treatments known to induce HSP, our findings could indicate that APP/PS1 neurons have attempted to adapt to reduce excitability (reduced baseline AIS) but are unable to do so to treatments that normally would induce HSP.

Cortical neurons in proximity to plaques were reported to have higher basal calcium levels in spines and dendrites (Kuchibhotla *et al*., 2008). Aβ oligomers of different varieties have been reported to bind various cell surface receptors such as PrP (Lauren *et al*., 2009), alpha7 nicotinic receptor (Sadigh-Eteghad *et al*., 2014) and Ephrins (Vargas *et al*., 2018), leading to an influx of calcium. Yet another hypothesis proposes that Aβ increases the cell membrane permeability for calcium (Kawahara & Kuroda, 2000; Kagan *et al*., 2002). To complicate matters, presenilins have also been implicated in the handling of Ca2+ stores independently of γ-secretase in AD transgenic mouse models (Lerdkrai *et al*., 2018). A recent study suggested that Aβ dimers could cause hyperactivity by inhibiting glutamate reuptake (Zott *et al*., 2019). Further, it was reported that Aβ oligomers can impair synaptic activity by repressing P/Q calcium channels (Nimmrich *et al*., 2008). While we prepared synthetic Aβ in DMSO, which prevents the formation of fibrils, Aβ forms amyloids with time in culture (Takahashi *et al*., *2004*) and progressively aggregates at synapses (Takahashi *et al*., 2002; Takahashi *et al*., 2004; Willén *et al*., 2017), consistent with Aβ aggregation at synaptic compartments(Bilousova *et al*., 2016). Moreover, while Aβ and APP influence synaptic activity, neuronal activity also regulates APP cleavage and Aβ generation (Kamenetz *et al*., 2003); increased neuronal activity can increase both the generation and degradation of Aβ (Kamenetz *et al*., 2003; Tampellini *et al*., 2009). Thus, converging data indicate that both APP and Aβ are important for regulating neuronal activity. Among questions that remain to be answered are which specific aspects of neuronal activity APP and Aβ regulate/influence. Many transgenic models of AD exhibit epileptic seizures and hyperactivity (Scharfman, 2012; Born *et al*., 2014), and even models overexpressing wild-type human APP develop seizures, which could be consistent with our data. Importantly, hyper-synchrony in AD transgenic mice could be rescued by genetic suppression of APP over-expression (Born *et al*., 2014). A better understanding of the neuron subtypes and molecular mechanisms involved in early Aβ/APP-induced hyperexcitability and synapse dysfunction might provide not only new insights into the disease, but also to new treatment strategies for AD.

## Acknowledgments

We thank Bodil Israelsson for technical assistance as well as master’s students Ainoa Pilkati and Mohammed Rahman for help with analyzing data. We appreciate the support of MultiPark, Hjärnfonden, Alzheimerfonden, Kockska stiftelsen, the Swedish Research Council (Grant #2019-01125), and the Olav Thon Foundation.

## Figure legends

**Supplemental figure 1. Live-cell imaging method for assaying spontaneous calcium transients. a-b)** Representative traces of spontaneous activity over a 2 min period in neurons from WT and APP/PS1 mice, respectively. **c-d)** Raster plots showing overall calcium transients (spikes represented as green dots) in WT and APP/PS1 neuron field of views (FOVs); ROI# or neuron number on the Y-axis and time in minutes on the X-axis.

**Supplemental figure 2. APP/PS1 neurons respond appropriately to both GABA inhibitors and tetrodotoxin. a)** Representative Raster plot of APP/PS1 neurons treated with vehicle (sterile milliQ water); neurons were imaged every 100 ms for 5 min with detected spikes represented as green dots. **b)** Representative raster plot of APP/PS1 neurons treated with 10 μM of picrotoxin. **c)** Representative raster plot of APP/PS1 neurons treated with 20 μM of bicuculline. **d)** Representative raster plot of APP/PS1 neurons treated with 1 uM of tetrodotoxin (TTX). **e)** Graph shows frequency from WT neurons treated with 1 μM TTX or 20 μM bicuculline. Note that TTX significantly reduces calcium oscillations whereas bicuculline significantly increases them compared to vehicle treated CTRL (TTX mean = 0.01429 CI = −0.01421-0.04278, n = 70; Vehicle mean = 2.923, CI = 2.094-3.752, n = 209; bicuculline mean = 25.00, CI = 23.04-26.97, n = 104, p values compared to vehicle: TTX = 0.0001 and bicuculline = 0.0001, Kruskal-Wallis test). **f)** Graph demonstrating acute effects of TTX and bicuculline in APP/PS1 neurons. Note that similar to in WT neurons, TTX blocks activity while bicuculline raises activity (TTX mean = 0.1644 CI −0.07614-0.4050, n = 118; vehicle mean = 5.164, CI = 4.140-6.188, n = 188; bicuculline mean = 13.77, CI = 10.49-17.06, n = 97, p values compared to vehicle: TTX = 0.0001 and bicuculline = 0.0379, Kruskal-Wallis test).

