## Supplementary figures and images for "Aβ/APP-induced hyperexcitability and dysregulation of homeostatic synaptic plasticity in models of Alzheimer’s disease"

### Supplemental figures

Supplementary figure 1

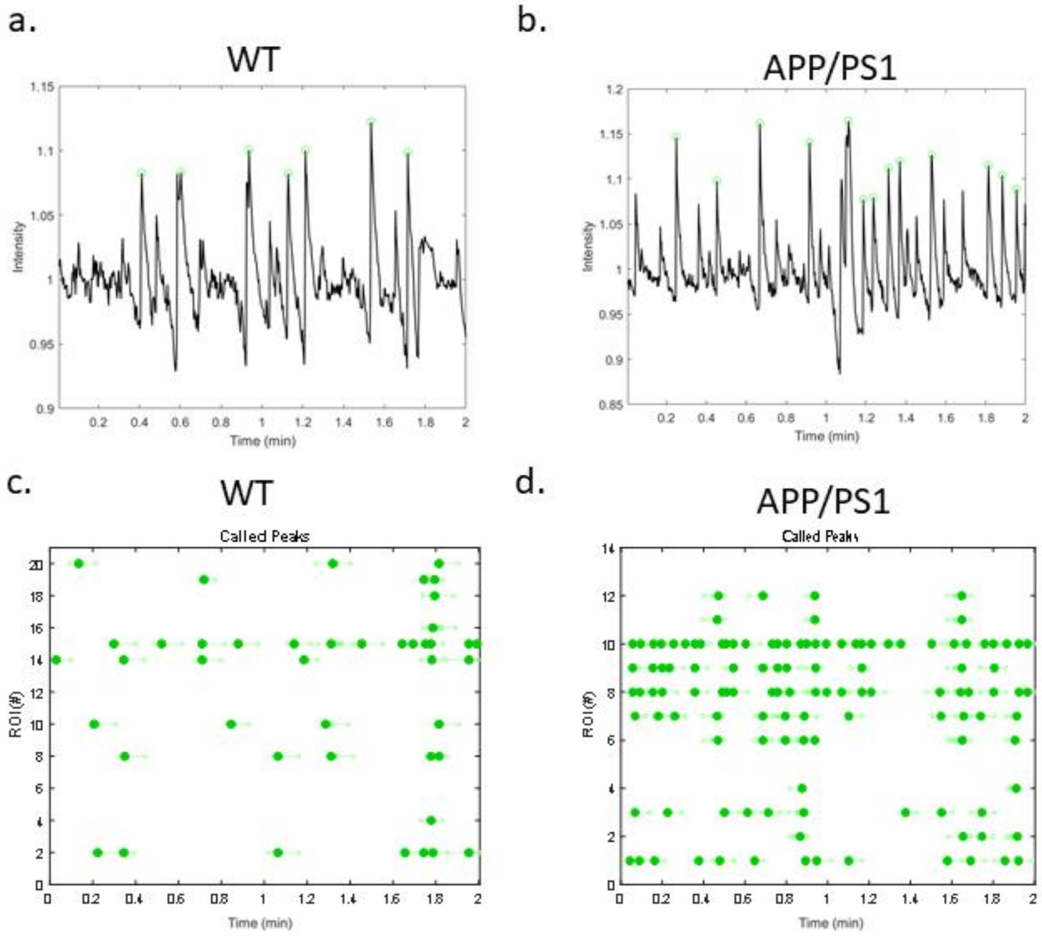

Supplementary figure 2

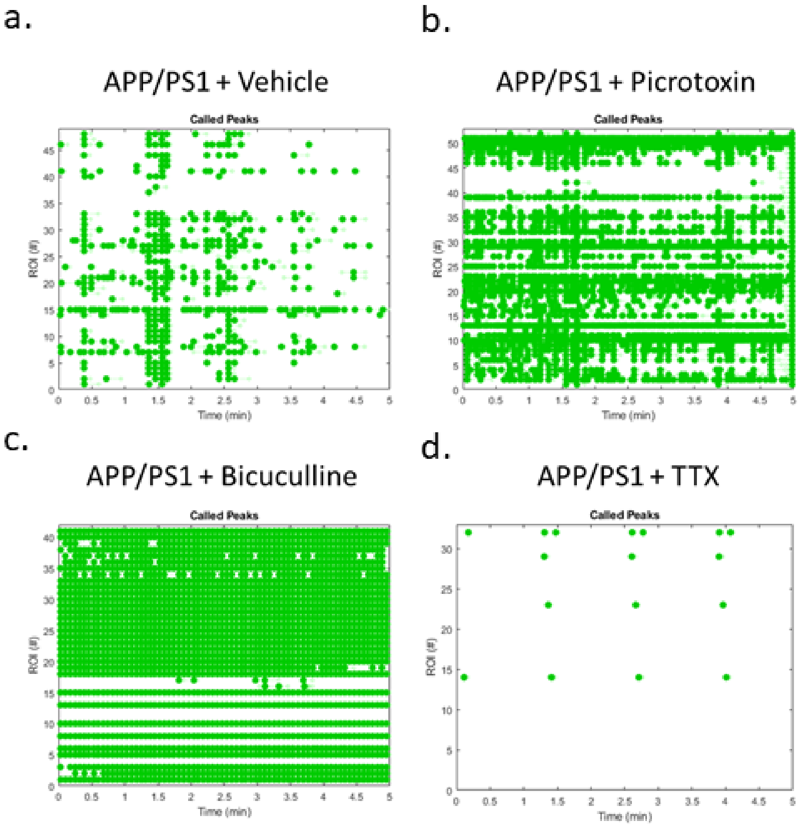


Acute treatment TTX or Bicuculline

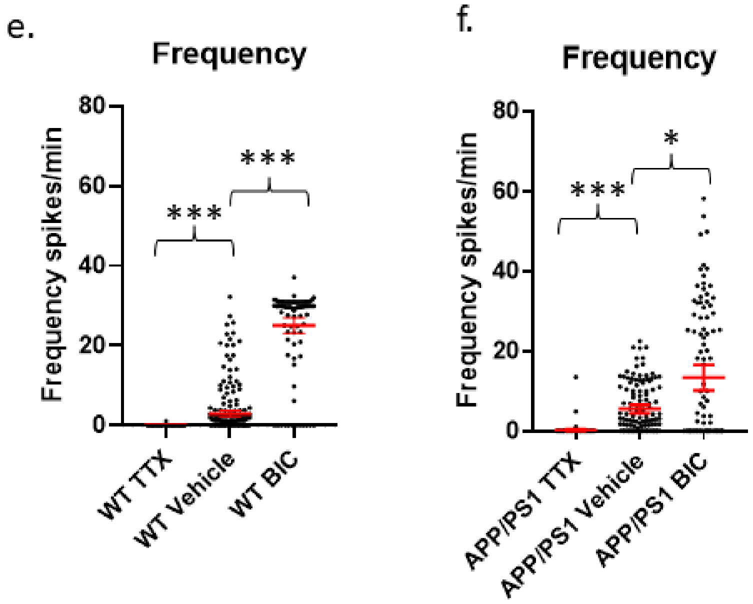
